# Compromised 2-start zigzag chromatin folding in immature mouse retina cells driven by irregularly spaced nucleosomes with short DNA linkers

**DOI:** 10.1101/2025.01.16.633430

**Authors:** Brianna Kable, Stephanie Portillo-Ledesma, Evgenya Y. Popova, Nathan Jentink, Matthew Swulius, Zilong Li, Tamar Schlick, Sergei A. Grigoryev

## Abstract

The formation of condensed heterochromatin is critical for establishing cell-specific transcriptional programs. To reveal structural transitions underlying heterochromatin formation in maturing mouse rod photoreceptors, we apply cryo-EM tomography, AI-assisted deep denoising, and molecular modeling. We find that chromatin isolated from immature retina cells contains many closely apposed nucleosomes with extremely short or absent nucleosome linkers, which are inconsistent with the typical two-start zigzag chromatin folding. In mature retina cells, the fraction of short-linker nucleosomes is much lower, supporting stronger chromatin compaction. By Cryo-EM-assisted nucleosome interaction capture we observe that chromatin in immature retina is enriched with i±1 interactions while chromatin in mature retina contains predominantly i±2 interactions typical of the two-start zigzag. By mesoscale modeling and computational simulation, we clarify that the unusually short linkers typical of immature retina are sufficient to inhibit the two-start zigzag and chromatin compaction by the interference of very short linkers with linker DNA stems. We propose that this short linker composition renders nucleosome arrays more open in immature retina and that, as the linker DNA length increases in mature retina, chromatin fibers become globally condensed via tight zigzag folding. This mechanism may be broadly utilized to introduce higher chromatin folding entropy for epigenomic plasticity.

## Introduction

In eukaryotic chromatin the DNA is discontinuously supercoiled to form “beads-on-a-string” chains of repeated 10-nm units (1–3) called nucleosomes (4). The nucleosome “beads”, each containing 145-147 bp of DNA making 1.7 left superhelical turns around an octamer of histones H2A, H2B, H3, and H4 in the nucleosome core (5,6) are connected by variable linker DNA “strings”. The linker DNA length varies dramatically amongst cells from different eukaryotic organisms and tissues (7,8) ranging from the 7 bp in fission yeast (9) to ∼100 bp in echinoderm sperm (10). The average core and linker DNA lengths combined comprise the nucleosome repeat length (NRL), which can be measured experimentally. Even within the same genome, the NRL in transcriptionally active chromatin domains may be up to 40 bp shorter than that in repressed or noncoding heterochromatin domains (11).

In living cells and at physiological conditions *in vitro*, nucleosome chains are packed into the higher-order structures (12,13). The nucleosome “beads-on-a-string” arrays (the primary structural level) fold longitudinally into the 30-nm chromatin fibers (the secondary structural level). These chromatin fibers then associate latitudinally to form large self-associated particles (tertiary structures) that may globally merge to form condensed interphase heterochromatin and metaphase chromosomes (14,15). The tertiary-level nucleosome condensates do not show any distinct secondary structures (16) and may form either liquid droplets or semi-solid hydrogels, depending on the experimental conditions and the epigenetic states (17,18).

The 30-nm fibers have been observed in the nuclei of some non-dividing cells with long NRLs. These include chicken erythrocytes (∼ 212 bp) and sperm cells of a starfish (∼ 222 bp) (19,20). Similar 30-nm fibers have also been reconstituted using arrays of nucleosome-positioning sequences with regular nucleosome linker lengths. Such reconstituted arrays showed a prominent two-start zigzag helical organization (21–27). However, despite many years of intense study, no regularly folded or helical structures above the 10-nm diameter were observed in the nuclei of proliferating cells (28–35).

Irregular chromatin folding may be caused by the intrinsic variability of the NRL. Indeed, experiments with chromatin fibers reconstituted with different NRLs showed strong effects of the nucleosome linker length on the chromatin fiber folding (10,36–40). Earlier modeling suggested that a mild stochastic variability of the NRL (+/3 bp) would significantly disorganize the 30-nm fiber making it to closely resemble the native chromatin (41). More recently, mesoscale chromatin models with alternating linker DNA lengths predicted polymorphic chromatin structures that form frequent loops and hairpin turns, thus dramatically deviating from the canonical chromatin fiber (42); non-uniform linkers representative of living systems were shown to enhance chromatin flexibility and encourage long-range contacts (43). Still, how variable the individual linkers are in the native chromatin species is largely unknown. This is due to the scarcity of direct measurements of individual linker length variability, and few structural or modeling studies based on the features of native chromatin rather than regular fibers with uniform NRLs and helical chromatin fibers.

The development of the mammalian eye retina provides an opportunity to study chromatin structural transitions in the process of cell maturation from proliferating retina progenitors to terminally differentiated non-dividing rod photoreceptors. The retina is a part of the central nervous system and has been studied extensively to analyze epigenetic and chromatin structure-mediated mechanisms of neuronal differentiation and maturation (44–46). Studies have also analyzed mechanisms of reprogramming cell differentiation that could lead to treatment of blindness and improving vision (47–50).

In mouse retinas, rod photoreceptors are the largest cell population (∼85%) and can be readily isolated in large numbers for biochemical experiments (>10^6^ cells per retina) (44). Heterochromatin in rod photoreceptors is accumulated in the middle of the nuclei and occupies over 70% of the nuclear diameter while the remaining euchromatin is located at the nuclear periphery (46,51). Though previous works have documented several factors controlling heterochromatin confluence in rod nuclei (50,52,53), the molecular and spatial aspects of chromatin higher-order folding underlying these epigenetic changes remain obscure. In particular, previous *in situ* transmission EM imaging of freeze-substituted rod photoreceptors followed by application of Fourier transform suggested the presence of 30-nm structures in the heterochromatin area but no distinct 30 nm chromatin fibers or other regular structures, helical or otherwise (51).

Previously we observed that the dramatic spatial segregation and chromatin condensation during retina development is accompanied by a notable increase in the NRL (45). Here, to examine the chromatin 3D structural transitions in the process of retina cell maturation, we apply Cryo-electron tomography (Cryo-ET) that can resolve biological structures embedded in thin layers of vitrified ice at nanoscale resolution (54,55). Unlike a single-particle reconstruction cryo-EM approach that can achieve higher resolution by averaging images of many thousand identical particles, Cryo-ET can resolve individual molecules and molecular assemblies, making it ideal for analysis of multiplex nucleosome chain conformations in chromatin (56). Cryo-ET was previously used to image short nucleosome arrays within thin sections of cryo-vitrified nuclei (57–59). With Cryo-ET analysis of partially unfolded long nucleosome arrays, we and others revealed a remarkable heterogeneity of linker DNA lengths (60,61). Here we report an intriguing relationship between linker length distribution and retina maturation: irregular and extremely short nucleosome spacings in immature retina progenitors compared to wider and more evenly spaced nucleosomes in mature counterparts. By *in situ* formaldehyde crosslinking coupled with Cryo-ET we found the impact of these compositional differences on 3D structure: chromatin in immature retina cells is enriched with i±1 interactions typical of open chromatin in transcribed genes, contrasting the predominately i±2 interactions in the mature retina, the latter typical of the two-start zig-zag folding. Finally, by computational mesoscale chromatin modeling, we show that the very short nucleosome linkers abundant in the immature retina cells strongly compromise the two-start zigzag formation and chromatin compaction by reducing the potential to form linker DNA stems. We propose that the increase of the nucleosome linker length is a key part of the molecular mechanisms driving heterochromatin spreading and gene repression during retina maturation and that a high frequency of short DNA linkers may be a general way of maintaining a more open chromatin to ensure high epigenetic plasticity in other immature progenitor cells.

## Materials and Methods

### Laboratory Animals

Wild-type C57Bl/6J (catalog #000664) mice were purchased from the Jackson laboratory and housed in a room with an ambient temperature of 25°C, 30–70% humidity, a 12 h light–dark cycle, and ad libitum access to rodent chow. This study was conducted using both male and female mice in accordance with the National Research Council’s Guide for the Care and Use of Laboratory Animals and all animal experiments were approved by the Penn State University College of Medicine Institutional Animal Care and Use Committee (protocol #00929).

### Retina Collection

Methods for retina isolation was performed as previously described (45). Whole eyes were collected from mice at the specified developmental stages. Retinas from whole eyes were dissected and placed into 1 mL of phosphate buffered saline (PBS) with 10-20 retinas collected per isolation.

### Isolation of retina nuclei and native chromatin

For standard nuclei isolation the collected retinas were triturated into a cell suspension by repeatedly pipetting up and down with a 1 mL plastic pipet tip. After tissue disassociation the cell suspension was centrifuged for 3 minutes at 500 g (4°C) in an Eppendorf 5810 R centrifuge. Cells were then resuspended in RSB buffer (10 mM NaCl, 3 mM MgCl_2_, 10 mM HEPES, pH=7.5) with 0.5% Igepal CA-630 (Sigma I3021), 1 mM PMSF, and protease inhibitor cocktail (Sigma P8849). Homogenization of cell suspension was carried out over 25 minutes with 30 strokes of a Dounce homogenizer with pestle B on ice then spun down for 7 minutes at 3,500g at 4°C in an Eppendorf 5810 R centrifuge with F-34-6-38 rotor. The resulting nuclei were then resuspended in RSB with 1 mM PMSF and protease inhibitor cocktail. For *in situ* formaldehyde fixation retinas were treated with formaldehyde to a final concentration of 0.2 – 0.4% prior to trituration for 15 minutes at room temperature with rotation. Following formaldehyde fixation glycine was added to a concentration of 125 mM and the mixture placed on ice for five minutes prior to centrifugation.

For micrococcal nuclease (MNase) digestion of native chromatin, MNase (Roche, #10107921001) was diluted to a concentration of 1.25 µg/mL in 1x bovine serum albumin (BSA) (New England Biolabs, #B9001S). For isolation of native chromatin nuclei were then treated with 2mM CaCl_2_ and incubated for five minutes in a 37°C water bath. Following incubation, MNase was added to a final concentration of 0.0125 µg/mL and incubated for 5 minutes a 37°C water bath with occasional shaking. To stop the enzymatic digestion, EDTA (Promega, #4231) was added to the nuclei suspension to final concentration of 5 mM and placed on ice for 5 minutes. The nuclei suspension was then spun down for 5 minutes at 10,000 rpm in in an Eppendorf 5810 R centrifuge with F-34-6-38 rotor cooled to 4°C. Following centrifugation, the pelleted nuclei were resuspended in cold TE buffer (10 mM Tris, 1 mM EDTA, pH=8.0) and incubated overnight at 4°C with end over end rotation. After incubation, the suspension was centrifuged again under the same conditions previously described. The soluble chromatin was collected in the resulting supernatant (S2). The final pellet, referred to as nuclear residue (NR), was resuspended in TE buffer. DNA concentration of the solubilized S2 or NR chromatin was measured by UV-vis spectroscopy. For assessment of DNA size following MNase digestion, aliquots of S2 chromatin were treated with SDS and proteinase K to final concentrations of 1% and 0.5mg/mL respectively and subjected to DNA electrophoresis. For subsequent biochemical analysis and imaging, the native chromatin and *in situ* fixed chromatin samples were subject to buffer exchange dialysis against HNE buffer (5 mM NaCl, 0.1 mM EDTA, 10 mM HEPES-NaOH, pH=7.5) for 24 hours in a 1:250 ratio in Slide-A-Lyzer™ MINI Dialysis Devices (Thermo Scientific, #69572).

### Cryo-electron microscopy and tomography

Retina chromatin samples with a concentration of about 0.2 mg/ml DNA were mixed with a suspension of 10 nm fiduciary gold particles (Sigma Aldrich cat.# 741957) treated by bovine serum albumin to prevent clustering. 3 μl chromatin samples were applied to Quantifoil R2/2 200 mesh copper grids (EMS Q250-CR2). Vitrification by plunging into liquid ethane was conducted using FEI Vitrobot Mk IV Grid Plunging System at 100% humidity, 4°C, and setting the blotting strength at 5, and blotting time at 3.5 sec. Cryo-EM imaging was conducted on Titan Krios G3i 300 kV electron microscope, equipped with a K3 direct electron detector (Gatan, CA) at the Penn State Hershey cryo-EM core as described (61). Tilt series were aligned using fiducials, CTF corrected and SIRT reconstructed using IMOD software suite (https://bio3d.colorado.edu/imod/).

### Regression tomogram denoising and 3D visualization

Regression denoising was accomplished in Dragonfly (ORS) using data synthesized by cryo-TomoSim (CTS, https://github.com/carsonpurnell/cryotomosim_CTS) as described (61,62). Regression denoised images were exported out as .tiff files and converted to MRC using the *mrc2tif* program within IMOD. The regression-denoised images were segmented into smaller subtomograms by IMOD/3dmod and inverted using “newstack” to generate subtomograms with positive intensity corresponding to high density. Data was visualized either with IMOD or Chimera. Codes for regression tomogram denoising (cryo-TomoSim) are publicly available at https://github.com/carsonpurnell/cryotomosim_CTS. Any additional information required to reanalyze the data reported in this paper is available from the corresponding author upon request.

### Stereological modeling and analysis

Stereological measurements were conducted as described before (61). Briefly, the SIRT-reconstructed tomograms were segmented into smaller subtomograms by IMOD/3dmod, inverted using “newstack”, and filtered by nonlinear anisotropic diffusion using IMOD command: “nad_eed_3d -n 30 -f -k 50” to reduce noise and enhance chromatin edges. The filtered subtomograms (both SIRT-reconstructed and regression-denoised) were exported into UCSF Chimera <u>(RBVI, Univ. San Francisco, CA)</u> for interactive visualization and analysis of nucleosome structures. In Chimera, the filtered volumes were fitted with nucleosome core X-ray crystal structure (pdb 2CV5 (63)) semi-automatically to correspondent electron densities in the volume using the ‘fitmap’ command and the nucleosomes were overlaid with centroids, center-to-center axes, and nucleosome planes using the structure analysis ‘Axes/Planes/Centroids’ tool. Each nucleosome in an array was numbered and the following measurements were recorded: center-to-center distance D to the next nucleosome (n+1) in the array; center-to-center distance N to the nearest nucleosome (n_x_) in the 3D space; angle *α* between the two axes connecting each nucleosome with the previous one (n-1) and the next one (n+1) in a chain: angle *β* between the planes of consecutive nucleosomes n and n+1; and angle *para* between the plane of each nucleosome (n) and the plane of the nearest nucleosome (n_x_) in the 3D space.

For a relatively minor fraction of nucleosomes (5.5% of total PN1 nucleosomes and 7.5% of total PN56 nucleosomes vitrified in HNE) the linkers were not resolved due to gaps in EM densities and corresponding distances D and angles *α* and *β* were excluded from the statistical analysis.

Open linker DNA exiting and entering consecutive fitted nucleosomes were manually traced with CHIMERA Volume Tracer tool using options: “place markers on high density” and “link consecutively selected markers”. Raw multi-segment open linker DNA linker lengths (o) were measured by Chimera “measure pathlength sel” command in nanometers and converted to the number of base pairs by assuming 0.34 nm per DNA base pair. The positions, at which the open DNA regions were exiting and entering the fitted 146 bp nucleosome cores were indicated as start (s) and end (e). Open DNA lengths (o) were measured between the start (s) and end (e) points at which naked DNA leaves the nucleosome cores. The distances of segments between the start (s) and end (e) points and the boundaries of the nucleosome core were subtracted from the nucleosome core DNA length (146 bp) resulting in the constrained core DNA length (C). The same segment distances were subtracted from the measured open DNA length (o) resulting in the linker DNA length (L).

The absolute nucleosome proximity values, (i±k) were obtained by subtracting the number of each nucleosome (n) from the number of its nearest nucleosome (n_x_) in the 3D space. The nucleosome contacts were defined as those occurring at distance N< 11 nm (double nucleosome disk radii) between the centroids as we did before (64). Nucleosomes involved in “trans” contacts were recorded when two nucleosomes from particles X and Y were in proximity < 11 nm, the linker connections were missing, and the unconnected nucleosomes in particles X and Y were at a distance of more than 28 nm (probability of physical connectivity at such distance is less than 1% from the analysis).

### Fluorescence microscopy

For fluorescence microscopy involving nuclear area measurements, isolated native and formaldehyde-crosslinked retina nuclei were resuspended in TE buffer (10 mM Tris pH 7.5, 1 mM EDTA) at ∼ 1:100 dilution and incubated at 4°C for 4 hr – overnight to allow for nuclei to expand. Wide-bore pipet tips were used to gently mix the nuclear suspension at 1:1 ratio with 0.2 μg/mL Hoechst 33258 in TE mixing gently. 0.5 ml aliquots of the nuclei stained with Hoechst 33258 were applied onto Poly-L-lysine glass slides (ThermoFisher 47100) in the areas encircled with hydrophobic Liquid Blocker pen, and incubated for 30 min in a chamber to settle down. The glass-attached nuclei were washed briefly with TE, then incubated in TE for 1 – 2 hr. Slides imaged by fluorescence microscopy using inverted Nikon Eclipse 2000 microscope and analyzed with NIS-Elements software. No coverslips were used to prevent bursting of unfixed nuclei while taking area measurements. Nuclei area measurements taken for ∼200 Hoechst-stained cells for each sample. Statistical analysis performed with Prism 10 using one-way ANOVA and Kruskal-Wallis test for multiple comparisons.

### High resolution DNA gel electrophoresis

DNA gel electrophoresis of in 1.1% agarose gel (Lonza, SeaKem LE Agarose, #50004) and Tris/acetic acid/EDTA (TAE) buffer (Bio-Rad, cat.# 1610743) was conducted as described (61). The gels were post-stained with GelRed (Biotium, #41003) and imaged digitally. For analysis of NRL the relative migration distance (Rf) of the DNA standards and sample DNA bands were measured using ImageJ software (65) (National Institutes of Health). A standard curve was generated by plotting the Rf of the DNA standards against the log(Molecular weight) of the DNA standards with Excel (Microsoft Excel for Office 365). The DNA length (bp) of each sample was then calculated with the resulting linear equation. To determine the NRL the DNA length was divided by the number of nucleosomes represented by the corresponding DNA band.

### SDS-PAGE

For verification of histone protein integrity native chromatin samples were dissolved in SDS-containing loading buffer and the electrophoresis was carried out in 18% acrylamide gels as described (66). Gels were post-stained in Brilliant blue R250 (FisherBiotech #FL-04-0598) and imaged digitally.

### Mg^++^-dependent self-association of chromatin

For in-gel chromatin self-association assays native chromatin samples in HNE buffer were mixed with increasing concentrations of MgCl_2_ (Sigma Aldrich, cat. # M1028) then incubated on ice for 20 minutes. Samples were then centrifuged at 12,000 rpm, 4°C, for 10 min in an Eppendorf 5810 R centrifuge with an F-34-6-38 rotor. The supernatants were collected and loading buffers containing 12% glycerol, 40 mM EDTA, 2% SDS or 8% glycerol, 10 mM EDTA, 1% SDS were added to the supernatant and pellet respectively. DNA samples were loaded by increasing concentration of MgCl_2_ onto a 1% agarose gel (Lonza, SeaKem LE Agarose, #50004) in TAE buffer for 40 minutes at 3V/cm then post stained with GelRed (Biotium, #41003). To determine the percentage of DNA remaining in either the supernatant or pellet the intensity of the DNA bands were quantified with ImageJ software (65). The percentage of DNA present in both the supernatant and pellet fractions in order of increasing concentrations of MgCl_2_ were entered into Prism version 10.0.0 for Windows (GraphPad Software) in order to interpolate a standard curve (Sigmoidal, 4PL, X is concentration). The resulting IC50 values for both the supernatant and pellet fractions were averaged to determine the concentration of MgCl_2_ at which 50% of the chromatin sample was precipitated.

For Cryo-ET imaging of magnesium induced condensation, chromatin samples in HNE buffer with a final concentration of 200 μg/mL were treated with MgCl_2_ solution and incubated on ice for 20 minutes prior to sample preparation.

### Genomic qPCR

For genomic quantitative real-time PCR (qPCR) was performed on CFX Connect Real-Time PCR System (Biorad) with iTaq Universal SYBR Green Supermix (Biorad, # 1725121) according to the manufacturer’s instructions.

### Quantification and statistical analysis

Violin plots were generated using Prism vs. 10.0.0 (Graph Pad). All other plots were generated in Excel (Microsoft). Average and standard deviation values were obtained from at least three tomograms and at least two independent biological samples. *p-*values representing probability associated with a Student’s two-sample unequal variance t-test with a two-tailed distribution are shown on the graphs; these values were calculated using Excel (Microsoft). Nonsignificant difference (ns) is shown for *p* > 0.05. Datasets with nonsignificant difference were additionally examined by nonparametric Kolmogorov-Smirnov test using Prism. In cases where the values are significant by Kolmogorov-Smirnov test but not by standard t-test (due to non-Gaussian distribution), the *p-*values resulting from the latter test are indicated by asterisks and are given in the corresponding figure legends. The numbers of nucleosomes (n) or nucleosome arrays (n’) accounted for in each test are given in the corresponding figure legends.

### Mesoscale Modeling of chromatin fibers

To further investigate changes in chromatin architecture during the maturation of retina cells, we apply our chromatin mesoscale model (reviewed in (67) and (68)) to simulate chromatin fibers typical of the PN1 and PN56 stages. To model the chromatin fibers, we set linker DNA length distributions obtained from the Cryo-EM images of individual chromatin arrays by equilibrium Monte Carlo (MC). We choose the two samples that show the largest differences between PN1 and PN56. For each stage, PN1 and PN56, we simulate 23-nucleosome arrays. Due to our model resolution of 8.8 bp for linker DNA beads, that translates to linker DNAs lengths of 0, 17.65, 26.47, 35.29, 44.12, 52.94, 61.76, 70.59, 79.41, and 114.71 bp. Both the PN1 and PN56 fibers are simulated with saturated linker histone (LH) of 1 molecule per nucleosome. To study the role of salt concentration on fiber architecture, each system is simulated with a NaCl concentration of 5 and 150 mM, and with the addition of 1mM Mg^2+^. The effect of 1mM Mg^2+^ is introduced implicitly by allowing the two linker DNAs to be closer and by reducing the DNA persistence length to 30 nm (69). Briefly, our chromatin mesoscale model combines coarse-grained representations of nucleosome cores, histone tails, linker DNA, and LHs within chromatin fiber arrays. Each chromatin element is coarse grained at a different level of resolution. The nucleosome cores are treated as disks with 300 Debye-Hückel discrete surface charges calculated by the DiSCO algorithm to mimic the electrostatic environment of the atomistic nucleosome (70). The N-terminal tails of histones H2A, H2B, H3, and H4, and the C-terminal tail of H2A are coarse grained as 5 residues per bead and attached to the nucleosome surface; each bead has a charge equal to the sum of the charges of the 5 residues that compose the bead (71). In total, there are 50 tail beads per core: 16 for H3, 10 for H4, 14 for H2A, and 10 for H2B. Similarly, LH is coarse grained as 28 beads with a resolution of 5 residues per bead, where 6 fixed beads describe the globular head (GH) and 22 flexible beads describe the long C-terminal domain (CTD) (72). The adjustment of charges on each LH bead for different concentrations of monovalent salt are calculated with the DiSCO algorithm. The linker DNA connecting nucleosomes is treated with a combined worm-like chain and bead model (73). The length of the segments connecting beads is *l_0_* = 3 nm, which produces a resolution of 8.8 bp based on *N_bp_ = l_0_/rise*, where rise is 0.34 nm, the distance between consecutive bps in a B-DNA. Each linker DNA bead has a salt-dependent charge calculated based on the Stigter procedure (74).

The energy function of the model includes stretching, bending, and twisting local terms for the linker DNA, stretching and bending terms for the tails and LHs, and electrostatic and excluded volume terms among each pair of beads as follows:

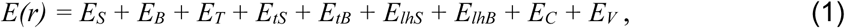

where ***r*** is the collective position vector. Details on each energy term can be found in the supporting information of Li et al. (75).

To obtain configurational ensembles in thermal equilibrium, we perform MC simulations of 20 copies of each PN1 and PN56 fibers started from different random seed and different residual twist DNA value of −12°, 0°, or +12°. Each copy is simulated for 40 million MC steps and the last 10 million steps of each trajectory are used for analysis.

For each system, we calculate the packing ratio as the number of nucleosomes in 11 nm of fiber, based on:

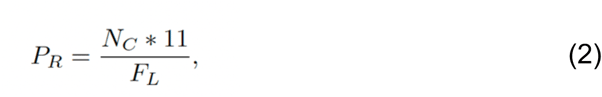

where *N_C_* is the number of nucleosomes and *F_L_* is the fiber length calculated by defining an axis that passes through the nucleosome cores. The fiber axis is defined as a three-dimensional parametric curve composed of three piece-wise polynomials created by performing a cubic smoothing spline interpolation (with smoothing parameter of 0.35) to the nucleosomes *x, y,* and *z* coordinates. The polynomials are evaluated at each nucleosome to obtain their center position in each direction. Then, the Euclidean distance between every nucleosome *i* and *i* + 2 across the fiber is calculated as the difference between the vectors defined by the center position in the *x, y,* and *z* directions. All Euclidean distances are then summed to obtain the total fiber length.

We also calculate the internucleosome interactions as follows. Two nucleosomes *i* and *j* are considered to be in contact if the distance between any element (core or histone tails) of nucleosome *i* and any element of nucleosome *j* is less than 2 nm. Nucleosome contact frequencies are calculated for the combined configurational ensemble that contains 2000 configurations, or 100 configurations from each of the 20 individual trajectories. The frequencies are then normalized by the total number of configurations. Contact matrices *I′(i, j)* are further decomposed into one-dimensional plots *I(k)* that depict the magnitude of *i, i ± k* interactions as follows:

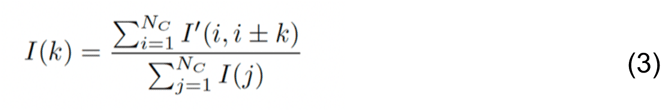

where NC is the total number of cores.

For the PN1 and PN56 fibers, both simulated at 5 and 150 mM NaCl, we create fan plots to represent, along each nucleosomal plane, the linker DNA cumulative and average positional distribution across a single trajectory. We determine the position vector of each DNA bead, **d**_ij_ , in the frame of reference of the parental nucleosome core with center of mass position *r_i_* and orientation {**a***_i_*, **b***_i_*, **c***_i_*}. The projected distribution in 3D is then denoted by **d***_ij_* = {**d**′_i*j,x*_, **d**′*_ij,y_*, d′*_ij,z_*}, where **d**′*_ij,x_* = **a***_i_* · (**d***_ij_* − **r***_ij_*), **d**′*_ij,y_* = **b***_i_* · (**d***_ij_* − **r***_ij_*), and **d**′*_ij,z_* = **c***_i_* · (**d***_ij_* − **r***_ij_*), and the projected distribution along the nucleosome plane is given by **d***_ij_* = {**d**′*_ij,x_*, d′*_ij,y_*}.

To assess quantitatively the degree of stem formation in the cores of PN1 and PN56, we measure the distances between the average positions of each pair of beads on the two linker DNAs associated with each core. That is, if any distance between a pair of corresponding beads located at the same position on the two linker DNAs i and j, i.e., i1 and j1, i2 and j2, and so on, is less than 2.5 nm, we consider that these beads contribute to a stem formation. Based on the total number of bead pairs contributing to the stem formation, we calculate a stem index that ranges from 0 to 1 through normalization. We perform this calculation using the average position of the DNA beads over a single trajectory of 40 million Monte Carlo steps, the same trajectory used for calculating the linker fan plots of Figure 7E.

For the PN1 and PN56 ensembles of 2000 chromatin configurations at 5 and 150 mM NaCl, we calculate the fraction of configurations in which each tail *t* (*t*=H2A–N-terminal, H2A–C-terminal, H2B–N-terminal, H3–N-terminal, and H4–N-terminal) is ‘in contact’ with another chromatin element *e* (*e*=core, DNA, or tails of a separate nucleosome).

We construct a two-dimensional matrix, where each matrix element T’ is defined as:

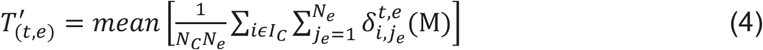

With 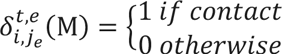

where NC is the total number of nucleosomes, Ne is the total number of chromatin elements, IC is a specific nucleosome along the chromatin fiber, and M is a specific chromatin configuration.

### We then normalize the interactions as

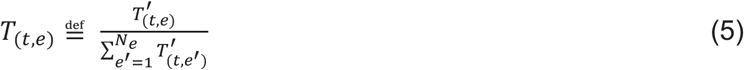

In each specific configuration M, we consider a t-kind tail of nucleosome *i* to be either free or in contact with only one Ne chromatin element based on the shortest distance between their beads and a cutoff of 2 nm.

## Results

### 1. Chromatin structural transitions during mouse retina maturation are driven by counterions and occur without major change in the protein composition

For chromatin isolation, we prepared cell nuclei from immature (PN1, postnatal day 1) and adult (PN56, postnatal day 56) mouse retina. Previously, using ultrathin tissue sections we observed that in PN1 retina, the chromatin is largely decondensed and the nuclei have a larger diameter than in PN56 retina, while the PN56 nuclei have a much smaller diameter (one of the smallest among vertebrate nuclei) with condensed and centrally positioned heterochromatin (45).

Isolated PN1 nuclei fixed by formaldehyde (Fig. 1 A) show uniform morphology with diameters significantly larger than PN56 nuclei with condensed chromatin. The isolated PN56 nuclei also show an almost uniform, small-diameter morphology and just one chromocenter (Fig. 1 B) contrasting with multiple chromocenters in PN1 nuclei (Fig. 1 A, D) consistent with previous in situ observations (45,46,51). Only about 10% of the PN56 nuclei had more than one chromocenter and larger diameter apparently due to the presence of other retina cells (Fig. 1 D). This result is consistent with rod photoreceptors being the predominant cell type in mature mouse retina (44,76) and demonstrates that we can efficiently isolate cell populations with either uniformly decondensed (PN1) or predominantly condensed (PN56) chromatin.

**Figure 1:**
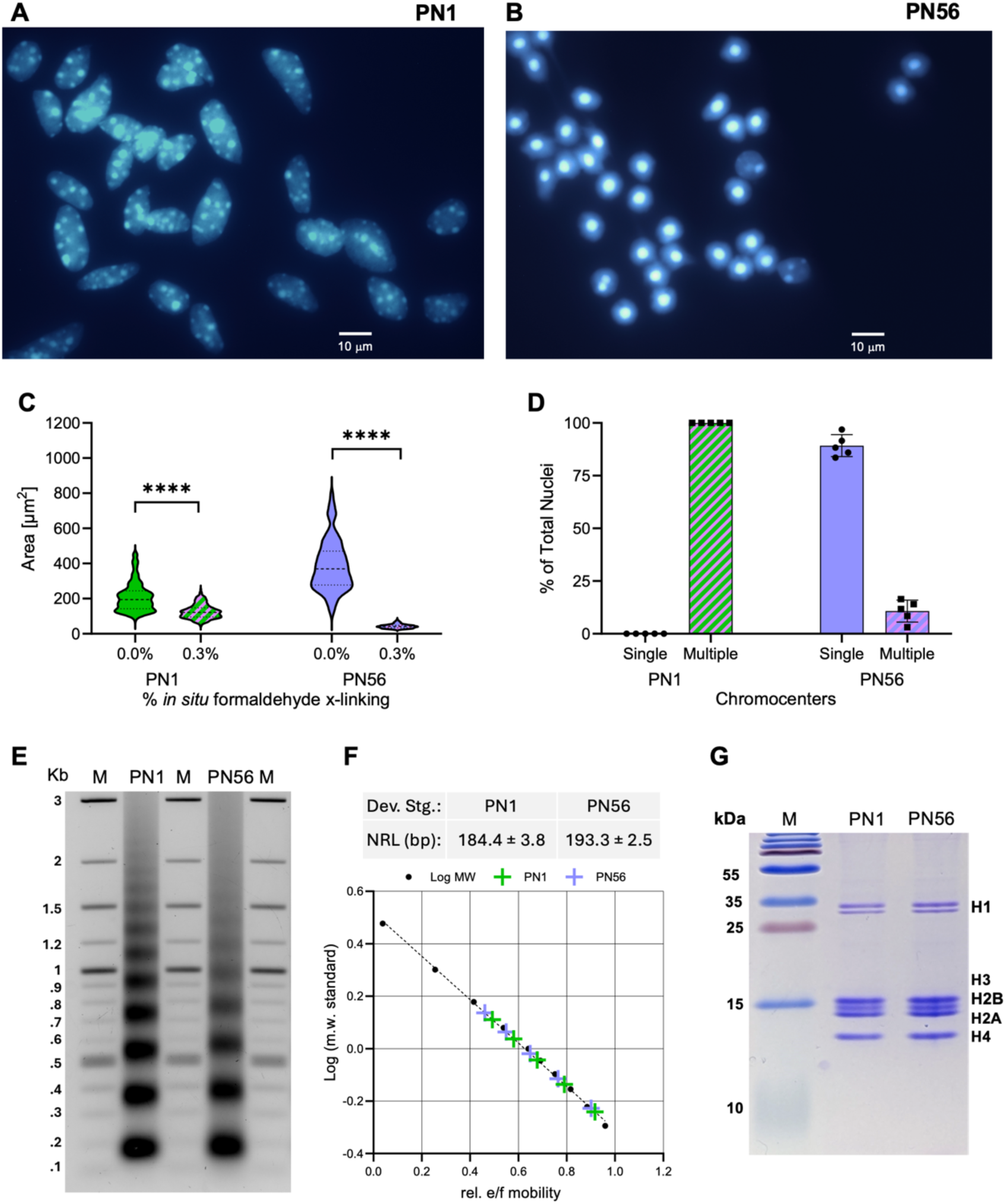
Isolation and characterization of the retina cell nuclei and soluble chromatin. A-B: Fluorescence microscopy imaging (Hoechst 33258 staining) of immature PN1 (A) and mature PN56 (B) nuclei isolated from retina cells crosslinked *in situ* with 0.4% of formaldehyde. C: Violin plots show average diameters of the intact and crosslinked PN1 and PN56 nuclei. D: Bar graphs show the number of Hoechst-positive chromocenters per nucleus. E: Agarose gel shows DNA size markers (lanes M), and DNA of native MNase-digested chromatin isolated from PN1 and PN56 mouse retina cells. F: Standard curve from the migration distance (Rf) and log(m.w.) of DNA size standards and nucleosome repeat lengths (±SD) determined for PN1 and PN56 mouse retina cells. G: 18% SDS-PAGE gel stained by Coomassie R250 shows molecular mass markers (lane M) and histones of PN1 and PN56 mouse retina chromatin.

To examine whether the compact heterochromatin is independent of intracellular milieu, we isolated PN1 and PN56 nuclei and recorded their diameter by fluorescence microscopy. Remarkably, after isolation and exposure to low ionic strength TE buffer the PN56 nuclei appeared to be considerably larger than PN1 nuclei. However, formaldehyde crosslinking of the nuclei before isolation reversed the trend making PN56 nuclei much smaller than PN1 (Fig. 1 C). It thus appears that chromatin in adult retina nuclei has a higher potential for salt-dependent contraction than in the immature retina nuclei.

We isolated MNase-fragmented soluble chromatin from PN1 and PN56 cells, adjusting the MNase digestion conditions to result in nucleosome arrays containing ∼ 12 nucleosomes per particle similar to the size of the nucleosome arrays used in most Cryo-EM studies of reconstituted chromatin compaction (21,25,26) and our recent Cryo-ET work with native human chromatin (61). By electrophoretic analysis of DNA in PN1 and PN56 cells, we observed a significant increase in the NRL from ∼ 184 to ∼193 bp (Fig. 1 E, F). By SDS-page we observed that both the PN1 and PN56 isolated chromatin contained mostly histones with very little nonhistone protein (Fig. 1 G) suggesting that the NRL may be a key variable affecting chromatin folding during retina rod cells maturation.

### 2. The potential for forming tertiary chromatin structure is not significantly different between chromatin in the immature and mature retina cells

We induced chromatin condensation by increasing the concentration of divalent cation, Mg^2+^, which promotes compaction of nucleosome arrays into zigzag fibers at 0.5 - 1 mM (23,26) close to the physiological range of free Mg^2+^ concentration (77). Chromatin isolated from both PN1 and PN56 retina cells showed a sharp self-association around 1.1 mM without a significant difference between the two chromatin samples (Fig. 2 A-C) and rather similar to the Mg^2+^-dependent self-association of reconstituted nucleosome arrays (16,78) suggesting that the potential for forming tertiary chromatin structure is not significantly changed during retina maturation.

**Fig. 2:**
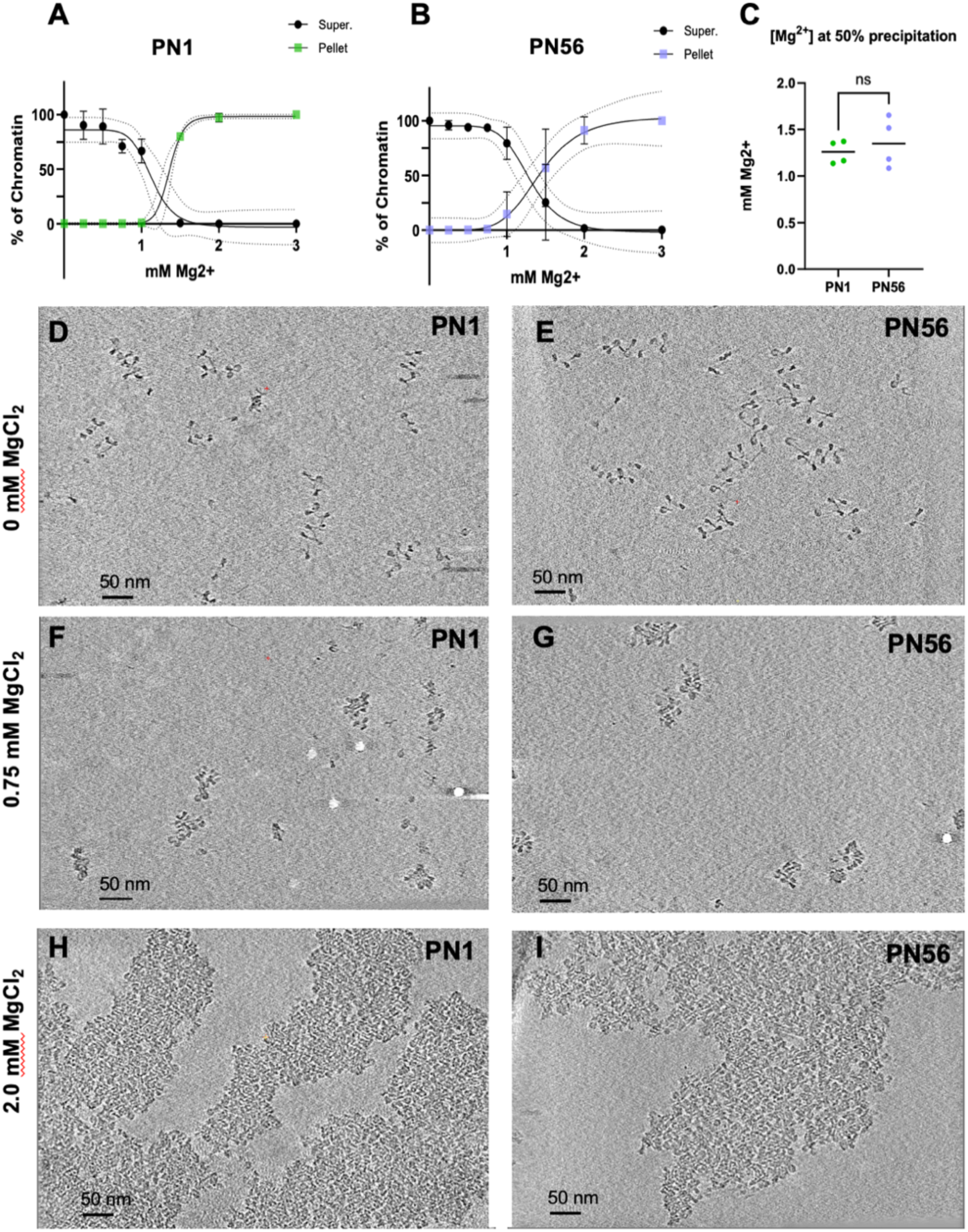
Cryo-ET imaging reveals tight nucleosome packing in Mg^2+^-condensed mouse retina chromatin. A-B: Graphs showing the % ratio of supernatant and precipitated PN1 and PN56 chromatin collected after mixing with the indicated concentrations of MgCl_2_. C: Mg^2+^-induced 50% precipitation points determined for the nucleosome arrays isolated from PN1 and PN56 cells. D-I: Representative Z-series slices from Cryo-ET tomograms of PN1 (D-F) and PN56 (G-I) chromatin embedded in vitrified ice at the indicated concentration of Mg^2+^ together with 10 nm fiduciary gold particles show open nucleosome arrays in HNE buffer without Mg^2+^ (D, G), compaction of individual particles at 0.75 mM Mg^2+^ (E, H)), and formation of bulky self-associates at 2.0 mM Mg^2+^(F, I). Mag: 53,000-x; scale bar 50 nm.

We then used Cryo-ET to examine native PN1 and PN56 chromatin vitrified in the low-salt HNE buffer (5 mM NaCl, 0.1 mM EDTA, 10 mM HEPES-NaOH, pH=7.5) with and without added Mg^2+^. The Cryo-ET tilt series were reconstructed by Simultaneous Iterative Reconstruction Technique (SIRT) using IMOD (79) software suite (https://bio3d.colorado.edu/imod/) and visualized as individual slices (Fig. 2 D-I).

With both PN1 and PN56 chromatin in the low-salt HNE buffer, we observed unfolded nucleosome arrays (Fig. 2 D, E) very similar to our previous Cryo-ET of native chromatin (61). In the presence of 0.75 mM Mg^2+^, the chromatin showed close juxtaposition of nucleosome disks without any helical zigzag fibers or other regular structures (Fig. 2 F, G). PN1 and PN56 chromatin vitrified at 2 mM Mg^2+^ showed prominent self-association in which nucleosomes merged to form bulky tertiary structures (Fig. 2 H, I). Though we could not resolve separate nucleosome arrays within the tertiary structures, the nucleosome packing showed apparent stacks of several nucleosomes (seen as edge-to edge) mostly observed on the structure periphery and internal regions where no stacking was detected. Thus, neither the biochemical characterization nor the overall “eagle-eye” Cryo-ET visualization of native retina chromatin show any dramatic chromatin structural changes in the process of retina maturation necessitating a more detailed ultrastructural analysis.

### 3. Cryo-ET and stereological analysis reveal significant changes of internucleosome spacing during retina maturation

To aid 3D visualization and resolve individual nucleosomes, we processed the tomograms of retina chromatin vitrified the low-salt buffer (Fig. 2 D, E) using deep learning-based regression denoised models as we recently described (61,62). This technique generates nearly noiseless tomograms, which facilitates the resolving of individual nucleosomes within most of the arrays detected in a tomogram (Fig. 3 A, E). Further, this approach enables improved visualization of the air/water interface so that the nucleosomes that could be damaged by the air contact are excluded from stereological analysis (61).

**Figure 3:**
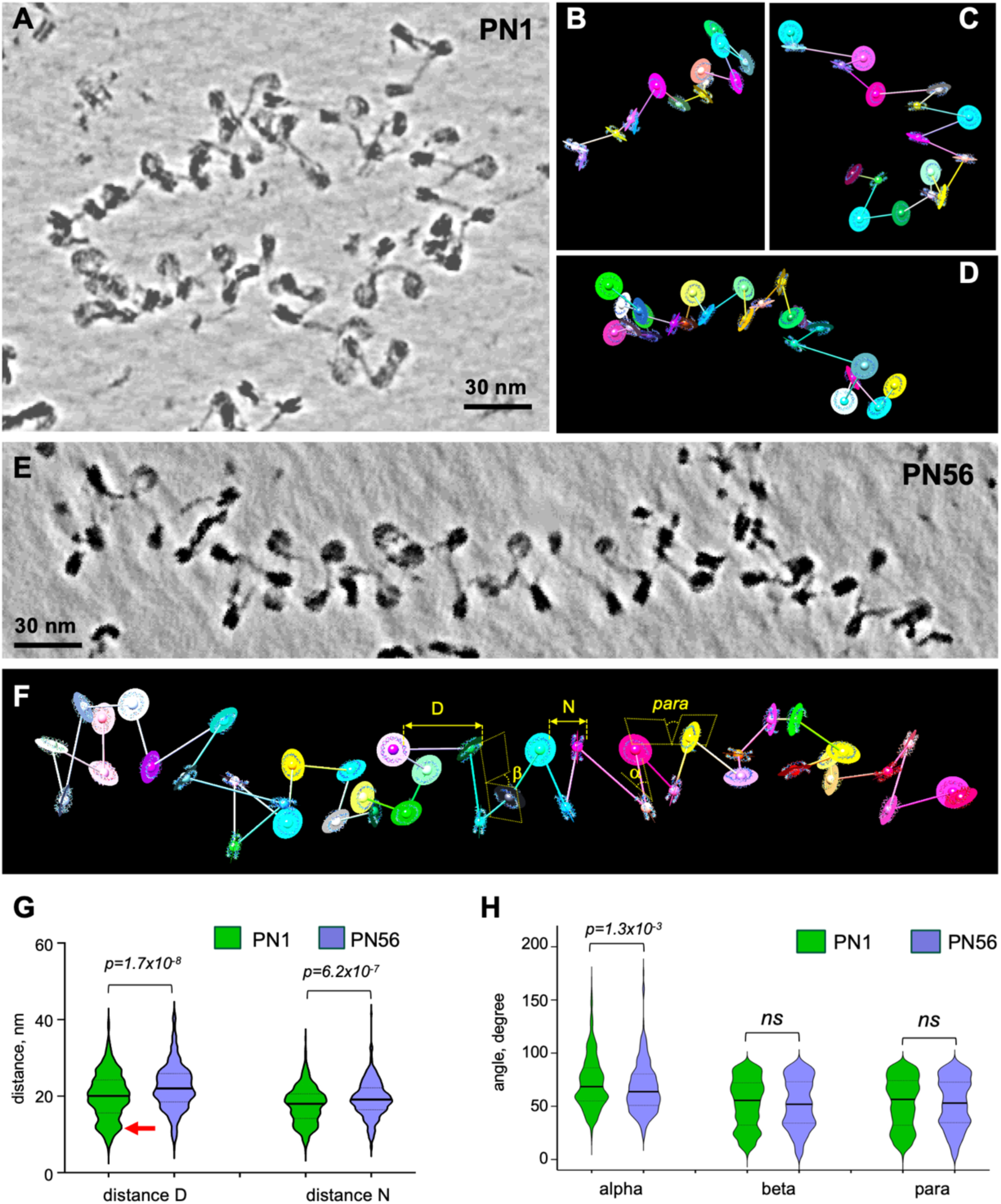
Cryo-ET and stereological modeling reveal significant changes of internucleosome spacing during retina maturation. A: Cryo-ET tilt series of the PN1 chromatin vitrified in low-salt HNE buffer (TS_1_1) and were tomographically reconstructed and processed by deep-learning denoising. A cropped image is shown as a composite of Z-series slices in IMOD. B-D: The cropped image of PN1 chromatin shown in (A) was processed to generate CAP models: B: CAP_1_5; C: CAP_1_3; D: (CAP 1_4). E: Cryo-ET tilt series of the PN56 chromatin in HNE buffer (TS_9_1) were tomographically reconstructed and processed by deep-learning denoising. A cropped image is shown as a composite of Z-series slices in IMOD. F: The cropped image of PN56 chromatin shown in (E) was processed to generate CAP model (CAP 9_2). The scheme shows variables measured from the CAP model: distances D and N and angles *α*, *β*, and *para*. G, H: Violin plots of distances D and N (F), and angles *α*, *β* and *para* (G) obtained for arrays of PN1 nucleosomes vitrified in HNE buffer (green shapes, n = 529 (D), 570 (N), 468 (*α*), 530 (*β*), 570 (*para*)) and arrays of PN56 nucleosomes vitrified in HNE buffer (violet shapes, n = 344 (D), 405 (N), 294 (*α*), 347 (*β*), 380 (*para*)).

When comparing the PN1 and PN56 nucleosome arrays, two features become readily apparent. First, PN1 contains nucleosome arrays with less even spacings than those in PN56. Second, PN1 arrays tend to easily change direction and shape in the 3D space while PN56 arrays tend to have more even zigzag organization and straighter array axis (cf. Fig. 3 A and E).

For quantitative stereological analysis, we cropped individual nucleosome array subvolumes from the SIRT-reconstructed cryotomograms, inverted the image to a negative, and processed them by nonlinear anisotropic diffusion filtering in IMOD. We then opened the filtered images in UCSF Chimera (https://www.cgl.ucsf.edu/chimera/ (80)) and built nucleosome chain models by rigid body fitting of the nucleosome core PDB models (PDB2CV5, (63)) into the density maps as we described (61). For each nucleosome we calculated the centroid and connected the centroids into a chain by axes (Fig. 3 B, C, D, and F) to generate centroid/axes/plane (CAP) models for each nucleosomal array. For each CAP model, we recorded five parameters for each nucleosome in PN1 and PN56 samples: 1) the center-to-center distance D from the next nucleosome in the chain, 2) the center-to-center distance N from the nearest nucleosome in 3D space, 3) the angle *α* between the two axes connecting each set of three consecutive nucleosomes in a chain, 4) the angle *β* between the planes of consecutive nucleosome pairs, and 5) the angle *para* between the planes of each nucleosome and its nearest neighbor in the 3D space (schematized on Fig 3 F). In addition, we also extracted individual array subvolumes from the corresponding deep learning-denoised cryotomograms, fitted them with nucleosomes, built CAP models, and conducted the same stereological measurements showing an excellent correspondence between the nonlinear anisotropic diffusion filtering and deep-learning regression denoising.

Due to the missing wedge in the tilt series (±60°) a minority of nucleosomes showed a gap in linker electron density so that 5.4% of all distances D in PN1 and 9.2% of all distances D in PN56 were not accounted for in the modeling. To test whether the nucleosomes with gaps in electron density might be structurally different from the total nucleosomes, we compared the total internucleosomal distances N and angles *para* (both recorded universally for all nucleosomes) with the fraction of those parameters connected without gaps (D^+^) and found no significant differences indicating an absence of nonrandom bias associated with the missing linkers.

Consistent with our MNase analysis showing about 10 bp increase in the linear length (Fig 1 G), the average distance D from one to the next nucleosome was significantly (p=1.7×10^-8^) increased from 20.0 to 22.3 nm during retina maturation (Fig. 3 G). Remarkably, in comparison to SD=±3 bp which accounts for the average DNA size in one MNase-produced band, the distribution of individual internucleosome distances was SD=±5.9 nm in both PN1 and PN56 corresponding to ±17.4 bp (ca. 0.34 nm per DNA base pair). This high nucleosome spacing heterogeneity is consistent with those previously observed in the metaphase HeLa chromosomes (60) and interphase K562 chromatin (61). Thus, the high heterogeneity in the nucleosome spacing is likely a general feature present in primary vertebrate tissues (such as mouse retina) as well as in proliferating cell cultures. Another striking feature in the analysis of D distribution (Fig. 3 G) is the notable peak of nucleosome distances at ∼ 11.5 nm in PN1 (red arrow) absent from PN56. The 11.5-nm peak in PN1 indicates prominent nucleosomes juxtaposed at a very short distance. The distributions of nearest neighbor distances N (which are not expected to be very different from D in the open chromatin) show the same tendencies with a significantly increased (p=6.2×10^-7^) N values in PN56 and the prominence of 11.5 nm nucleosome distance in PN1 (Fig. 3 G). In addition to PN1 and PN56, our stereological analysis of nucleosomes at postnatal day 21 (PN21) shows intermediate values between PN1 and PN56 samples, confirming that internucleosome distances gradually change during retina development.

In comparison to the significant changes in the internucleosomal distances, the changes in angles *α*, *β*, and *para* are rather modest (Fig. 3 H). Only angle *α* showed a significant (p=1.3×10^-3^) decrease from 72.9 to 67.1 degree; this change reflects an increase in linker histone H1 during retina maturation (45) which makes the angle between DNA linkers entering and exiting the nucleosome narrower and promotes formation of linker DNA stem (21).

### 4. Linker DNA tracing reveals nucleosomes with exceptionally short linkers and a wide variation of nucleosome linker length distribution in immature retina chromatin

While the above stereological analysis accurately measures the internucleosome distances and angles based on automatic fitting of the nucleosomes to the density maps, it does not allow one to measure the length of individual linkers that may be differentially folded or peeled of the nucleosome and thus conceal the actual length of DNA connecting the nucleosomes.

To further investigate the mechanism generating the surprisingly wide variation of internucleosome distances, we estimated the DNA linker length distribution by fitting nucleosome cores into deep-denoised images of open nucleosome arrays (Fig. 4 A), manually tracing the open DNA regions (o) between the start (s) and end (e) points where naked DNA leaves the nucleosome cores using Chimera Volume Tracer tool (Fig. 4 B), and subtracting the unwrapped nucleosome core DNA segments that did not fit into the observed core nucleosome volume resulting in the measured constrained core DNA length C (Fig. 4 C) and linker DNA length L (Fig. 4 D).

**Figure 4:**
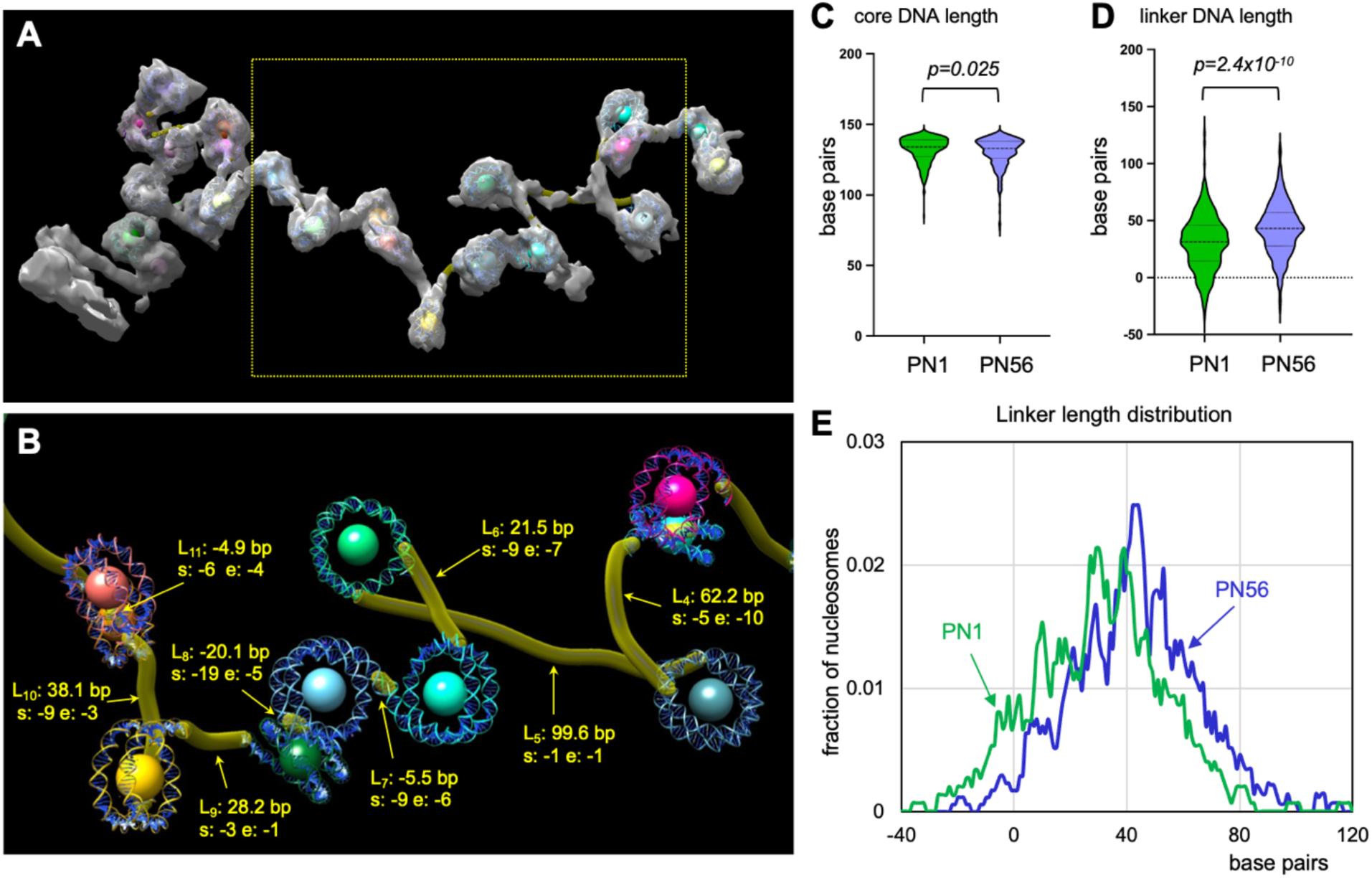
Tracing and comparison of linker DNA length in the PN1 and PN56 chromatin. A: A cryotomogram of the PN1 chromatin vitrified in HNE buffer (TS_1_1) was processed by deep-learning denoising, transferred to Chimera to generate CAP_1_4, volume-fitted with nucleosome core structures (pdb 2CV5), and shown as an isosurface. B: Open linker DNA in CAP_1_4 was manually traced with CHIMERA Volume Tracer tool. When the open DNA regions were exiting and entering the fitted nucleosomes at certain position within the 146 bp nucleosome core indicated as start (s) end (e), the unpeeled nucleosome core DNA base-pairs that did not fit into the traced volume were subtracted to give the linker DNA length (L) value. L4 (linker 3) to L11 (linker 11) indicated by arrows show the measured linker lengths (L) and the positions of the open DNA start (s) and end (e) in DNA base pairs. C: Violin plots show distributions of unpeeled core DNA lengths calculated for PN1 (green, n = 300) and PN56 (violet, n = 371) nucleosomes. D: Violin plots show distributions of linker DNA lengths L calculated for PN1 (green, n = 298) and PN56 (violet, n = 378) nucleosomes. E: Frequency distribution profile of linker length L calculated for PN1 (green) and PN56 (violet) nucleosomes.

Comparison of the core length DNA within the PN1 and PN56 shows only a minor difference (1.7 bp) that cannot explain the observed difference in nucleosome spacing. At the same time, L is increased by 11.9 bp in PN56 compared to PN1, showing that the nucleosome spacing increase during retina maturation is mostly due to the increased linker DNA length. When added to the number of the base pairs in the fitted nucleosome core PDB2CV5 (146 bp), the resulting NRL values (176.7 ± 24.1 bp for PN1 and 188.7 ± 22.6 bp for PN56) are rather close to those observed by agarose gel analysis of MNase-digested PN1 and PN56 chromatin (184 ± 3.8 bp and 193 ± 2.6 bp respectively, Fig. 1G) while the standard deviations in the distance between individual nucleosomes resolved by cryo-ET are dramatically higher than between the DNA gel measurements of averaged NRLs.

As with distance D, the measured DNA linker lengths (L) distribution showed a notable increase in the frequency of very short (∼10 bp) and negative length (<0 bp) linkers in PN1 (Fig. 4 E). The negative length linkers form when two cores closely overlap such as L_7_, L_8_, and L_11_ on Fig. 4 B and comprise 10.7% of all linkers in PN1. The very short linkers are found in the same arrays with the very long ones (e.g. L_4_ and L_5_) thus generating a very strong linker length heterogeneity in PN1. This linker length heterogeneity is also present in PN56, where though the negative length linkers are greatly diminished (3.2% of all linkers have negative length), the very short linkers and the long linkers co-exist in the same array. (This demonstrates that during the process of retina maturation and exit from cell cycle major changes in the primary nucleosome chain organization occur. These changes involve not only an increase in the average nucleosome linker length but also a striking decrease in the nucleosomes juxtaposed at a very short distance.

### 5. Chromatin from mature retina rod photoreceptors undergoes a stronger cation-driven compaction

Recently we observed that Mg^2+^-dependent condensation of native human chromatin led to the formation of prominent nucleosome disk stacking within the heteromorphic secondary structures that deviated from the regular 30 nm fibers (60,61). Since we observed a higher degree of heterogeneity in mouse retina chromatin than in human cells, we asked whether this chromatin still has a potential for cation-dependent condensation. Therefore, we induced PN1 and PN56 chromatin condensation by adding 0.75 mM Mg^2+^ to achieve maximal compaction without inducing self-association (Fig. 2), extracted sub-tomograms (Fig. 5 A, D), built CAP models (Fig. 5 B, C, E, F) and examined next nucleosome distances D, nearest nucleosome distances N, and the nucleosome plane angles *α*, *β*, and *para* (Fig. 5 G-J) as we did before for open chromatin (Fig. 3).

**Figure 5:**
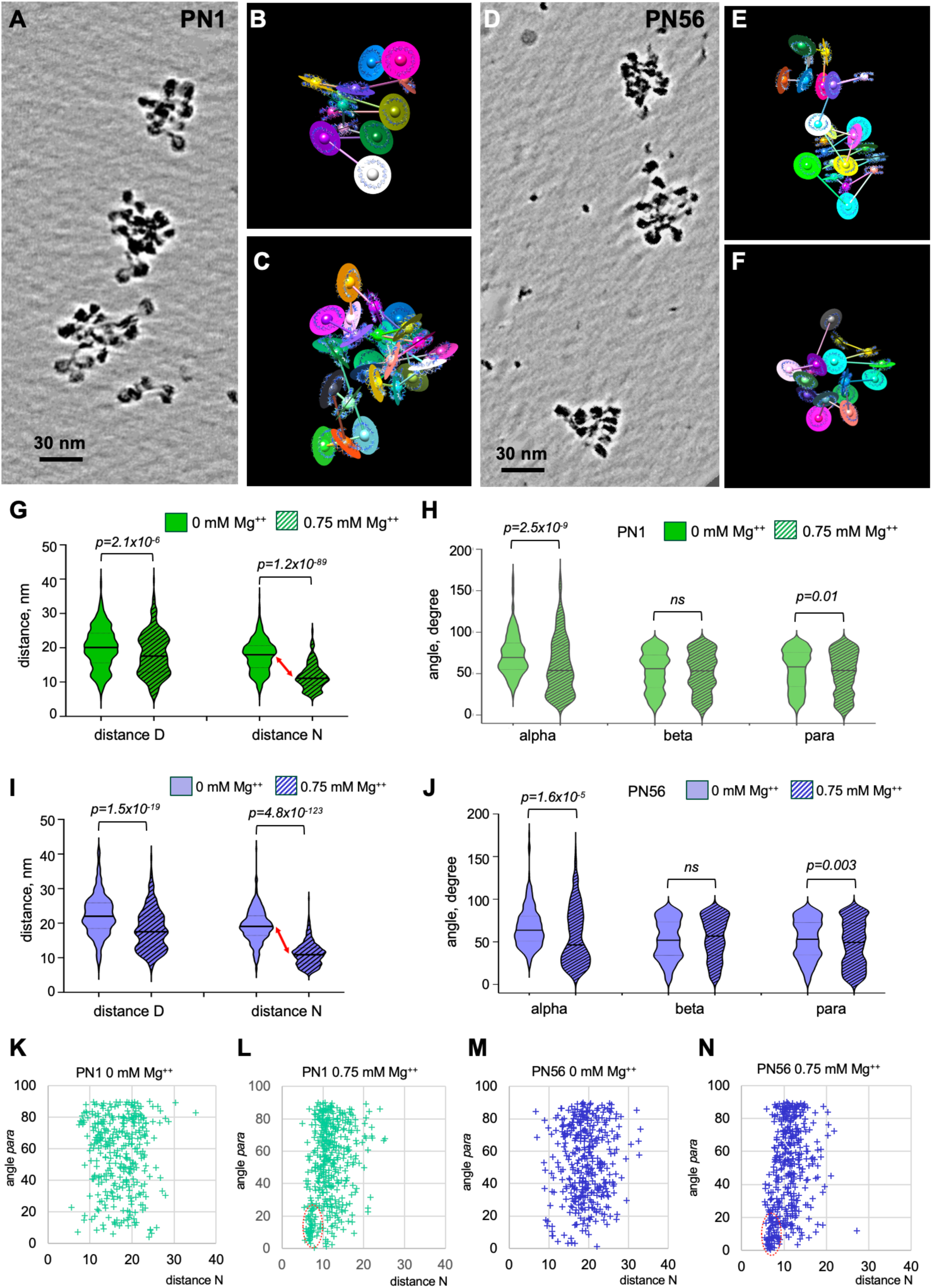
Cryo-ET and stereological analysis of immature and mature retina chromatin folding. A: Cryo-ET tilt series of the PN1 chromatin vitrified at 0.75 mM Mg^2+^(TS_3_2) were processed by deep-learning denoising and shown as a composite of Z-series slices in IMOD. B, C: cropped images of the PN1 chromatin shown in (A) were processed to generate CAP models CAP_3_11 (B) and CAP_3_10 (C). D: Cryo-ET tilt series of the PN56 chromatin vitrified at 0.75 mM Mg^2+^(TS_11_2) were processed by deep-learning denoising and shown as a composite of Z-series slices in IMOD. E, F: cropped images of the PN56 chromatin shown in (D) were processed to generate CAP models CAP_11_15 (E) and CAP_11_14 (F). G, H: Violin plots of distances D and N (G), and angles *α*, *β* and *para* (H) obtained for arrays of PN1 nucleosomes vitrified at 0 mM Mg^2+^ (empty green shapes, n = 529 (D), 570 (N), 468 (*α*), 530 (*β*), 570 (*para*) and arrays of nucleosomes vitrified at 0.75 mM Mg^2+^ (crossed green shapes, n = 334 (D), 431 (N), 284 (*α*), 345 (*β*), 430 (*para*)). I, J: Violin plots of distances D and N (G), and angles *α*, *β* and *para* (H) obtained for arrays of PN56 nucleosomes vitrified at 0 mM Mg^2+^ (empty violet shapes, n = 344 (D), 380 (N), 294 (*α*), 347 (*β*), 380 (*para*) and arrays of nucleosomes vitrified at 0.75 mM Mg^2+^ (crossed violet shapes, n = 310 (D), 399 (N), 249 (*α*), 312 (*β*), 399 (*para*)). K-N: Two-dimensional plots showing distributions of the internucleosomal angle *para* vs. the distance N for PN1 nucleosomes vitrified at 0 mM Mg^2+^ (K) and 0.75 mM Mg^2+^ (L) and PN56 nucleosomes vitrified at 0 mM Mg^2+^ (M) and 0.75 mM Mg^2+^ (N).

Comparative analysis of multiple nucleosomes vitrified at 0 mM Mg^2+^ and 0.75 mM Mg^2+^ in PN1 and PN56 chromatin showed that the distance between consecutive nucleosomes (D) was reduced rather modestly, by 10.0% of the original value for PN1 (Fig. 5 G) and 9.3% for PN56 (Fig. 5 I), consistent with relatively small linker DNA contraction upon condensation and similar to what we observed for human chromatin. In comparison, the average distance N between the nearest neighbor nucleosomes was reduced much more profoundly, by 34.4 % for PN1 and 42.7% for PN56 (Fig. 5 G, I), producing a prominent peak at ∼12 nm and a broader peak at ∼ 6.5 nm that correspond to the average distance between the perpendicularly proximal nucleosome disks and parallel stacked nucleosomes respectively as previously described (61). The overall degree of compaction by Mg^2+^ that we monitored by the average distance N between nearest nucleosomes was very close for the human (10.9 nm) and the PN56 mouse chromatin (11.0 nm) with notably less compact PN1 chromatin (11.6 nm).

Distribution of angle α shows a sharp and significant increase in average values for both PN1 and PN56 chromatin, generating a broad peak at ∼20° (Fig. 5 H, J). The angle *α*distributions are similar to that found in compact K562 chromatin but much wider than in compact clone 601 reconstitutes (61). This observation is consistent with heterogeneous angle α populations and likely reflects a variable association with linker histone H1 that reduces the angles between linker DNA (81,82). The angle *β* distributions are very wide in both PN1 and PN56 chromatin and do not change significantly upon chromatin compaction (Fig.5 H, J), indicating that the variable nucleosome plane orientations resulting from linker DNA length variations is not much altered during compaction consistent with the highly heteromorphic chromatin fibers observed by Cryo-ET for both immature and mature retina chromatin (Fig. 5 A-F).

In the both types of condensed retina chromatin, the frequency of internucleosomal angle *para* displayed a notable increase at values below 20° consistent with appearance of a fraction of stacked nucleosomes with near-parallel orientation of nucleosome discs. To reveal associations between the angle *para* and the internucleosomal distance N, we plotted their distributions on two-dimensional plots (Fig. 5 K-N). These plots show distinct areas below 25° and centered at the distance N ∼ 6.5 nm prominent at 0.75 mM Mg^2+^ (dashed ovals on Fig. 5 L, N) that are absent at 0 mM Mg^2+^ (Fig. 5 K, M). These areas correspond to the tightly stacked nucleosomes defined as a nucleosome separated from another nucleosome at distance N < 8 nm with angle *para* < 25° (61). Interestingly, with both PN1 and PN56 retina samples, the number of stacked nucleosomes, 10.7 and 12.5% respectively, appeared to be about 3-fold lower than the number of stacked nucleosomes (34.3%) in human K562 chromatin (61).

Thus, chromatin from mature PN56 mouse retina undergoes a greater degree of compaction than the immature PN1 retina chromatin, while maintaining high levels of secondary structure heterogeneity that are reflective of the intrinsic variability of the nucleosome positions in its primary structure.

### 6. In-situ crosslinking and characterization of soluble chromatin by Cryo-EMANIC

To reveal the features of chromatin architecture specific to the immature (PN1) and mature (PN56) mouse retina chromatin, we combined our EM-assisted nucleosome interaction capture (EMANIC) (64,69) with cryo-ET (Cryo-EMANIC) which involves crosslinking living cells with formaldehyde, followed by fragmentation of the nuclear chromatin by micrococcal nuclease, isolation and unfolding of chromatin fragments, and scoring nearest-neighbor nucleosome interactions by Cryo-ET (Fig. 6A). Unlike the previous transmission EM-based approach, Cryo-ET imaging is free from artifacts caused by attachment to the EM grid and heavy metal staining, which results in prominent “husks” partially coating crosslinked chromatin. This allows the quantification of nucleosome interactions with much lower background and without any protease treatment to open chromatin as was used for TEM analysis (64).

**Figure 6:**
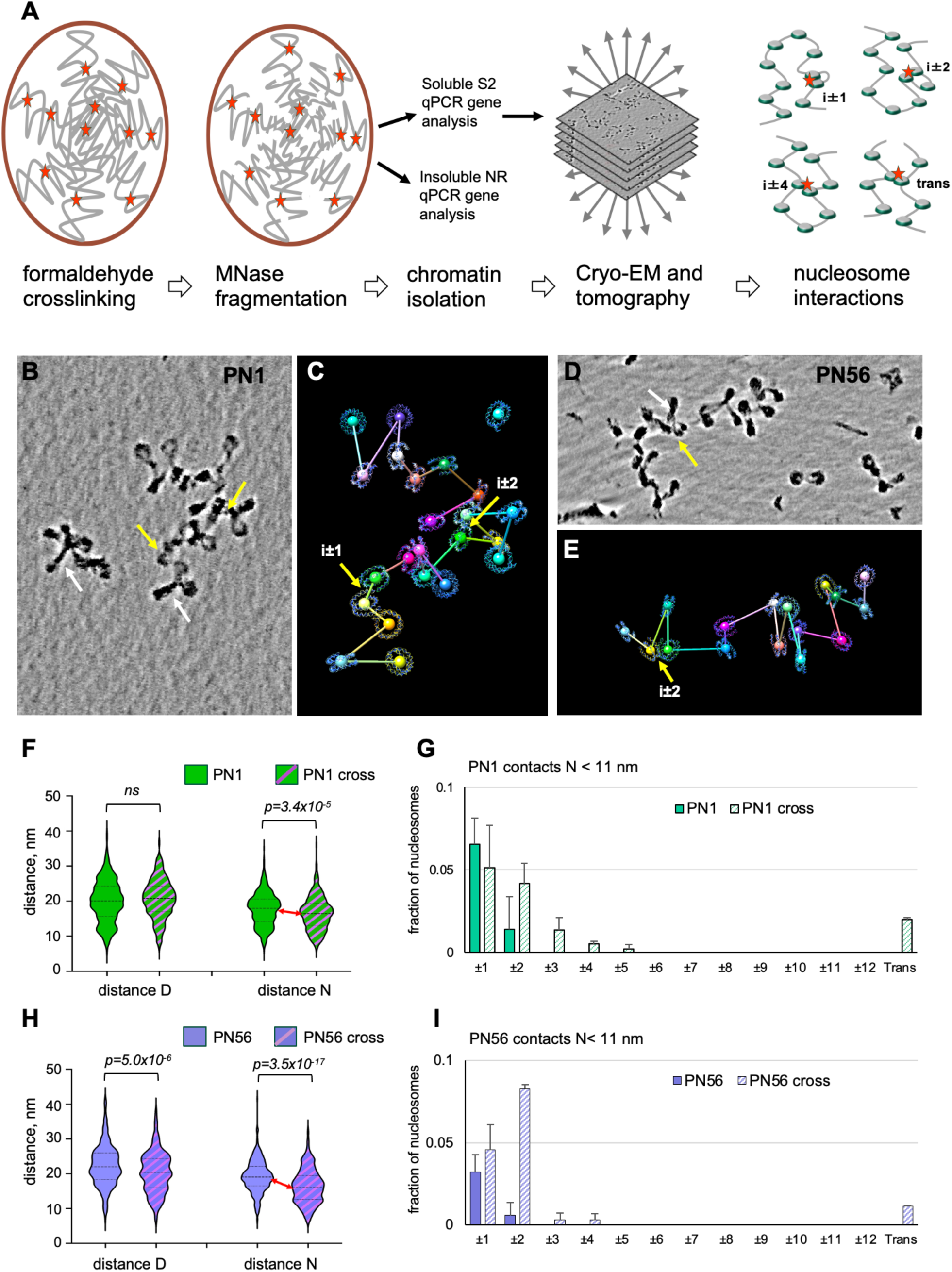
Cryo-EMANIC and analysis of nucleosome contacts *in situ*. A: scheme of the experimental procedure steps for Cryo-EMANIC. i±1, i±2, i±4 result from intra-fiber internucleosomal crosslinking; trans results from inter-fiber nucleosome crosslinks. B-E: Examples of cropped subtomograms of the *in-situ* crosslinked PN1 (TSS_8_1: B, C) and PN56 (TSS_15_2: D, E) chromatin vitrified in the low-salt HNE buffer processed by deep-learning denoising and shown as a composite of Z-series slices in IMOD (B, D) and used to generate CAP models CAP_7_4 (D) and CAP_15_8 (E). Planes are not shown to emphasize interactions. Yellow arrows indicate specific juxtaposed nucleosome cores (N < 11 nm) resulting from crosslinking. White arrows show nucleosome stems. F: Violin plots of distances D and N obtained for native PN1 nucleosome arrays (empty green shapes (n = 529 (D), 570 (N)) and *in-situ* crosslinked nucleosome arrays (crossed green shapes, n = 395 (D), 446 (N)). G: Distribution of nucleosome contacts at N <11 nm determined for native PN1 nucleosome arrays vitrified at 0 mM Mg^2+^ (green columns) as a fraction of total nucleosomes (n = 527) and *in-situ* crosslinked nucleosome arrays (crossed columns) as a fraction of total nucleosomes (n = 397). Error bars represent SD values. H: Violin plots of distances D and N obtained for native PN56 nucleosome arrays (empty violet shapes (n = 344 (D), 380 (N)) and *in-situ* crosslinked nucleosome arrays (crossed violet shapes, n = 350 (D), 416 (N)). I: Distribution of nucleosome contacts at N <11 nm determined for native PN56 nucleosome arrays vitrified at 0 mM Mg^2+^ (violet columns) as a fraction of total nucleosomes (n = 339) and *in-situ* crosslinked nucleosome arrays (crossed columns) as a fraction of total nucleosomes (n = 350). Error bars represent SD values.

To monitor the extent of formaldehyde crosslinking, we isolated PN1 and PN56 nuclei after a series of retina cell treatments with formaldehyde from 0 to 0.4%, transferred the nuclei into a low-salt HNE buffer that causes a dramatic unfolding of the unfixed nuclei and observed nuclei by fluorescence microscopy. Both PN1 and PN56 nuclei were efficiently crosslinked with 0.3 % formaldehyde causing an almost complete reduction of the nuclear area. We also observed that crosslinking with up to 0.2% formaldehyde did not inhibit the release of soluble chromatin; with more than 0.3% formaldehyde, we observed a significant decrease in solubility and therefore used crosslinking with 0.3% formaldehyde that released ∼73% soluble chromatin. We also monitored the gene composition of the soluble and insoluble chromatin fractions by genomic qPCR to observe that while some actively transcribed genes were underrepresented in the solubilized chromatin, the genes that represented the repressed and condensed chromatin were readily solubilized after 0.3% formaldehyde crosslinking.

Typical Cryo-ET images of nucleosome arrays isolated from crosslinked PN1 and PN56 nuclei are shown in Fig. 6, B, D. We processed the cryo-tomograms by deep denoising and conducted stereological analysis as before. For each nucleosome we calculated the centroid and connected the consecutive centroids by axes. For both PN1 and PN56 chromatin, the *in-situ* formaldehyde crosslinking resulted in a visible increase in the number of closely juxtaposed nucleosomes (yellow arrows on Fig. 6 B-E) and nucleosome linker “stems” (white arrows) as well as a significant decrease in the internucleosomal distances N (Fig. 6 F, H). The distance N was more prominently reduced for PN56 (by 14.7%) than for PN1 (7.0%) apparently reflecting the more condensed chromatin state in the mature retina cells.

For quantification, we defined direct nucleosome contacts as those occurring at distance N< 11 nm (double nucleosome disk radii) between the centroids as we did before (64) and determined relative nucleosome positions in the chain (i±k) between the contacting nucleosomes. Fig. 6 G, I show the relative abundance of cases where two consecutive nucleosomes are contacting (i±1), cases involving contacts (loops) over one (i±2) or more (i± >2) nucleosomes separating the crosslinked pair, and cases where two nucleosomes are contacting in-trans, between different arrays. Between the control and formaldehyde-crosslinking samples, the extent of the i±2 interactions increased most substantially: ∼3-fold for PN1 and ∼14-fold for PN56 (Fig. 6 G, I). At the same time, PN56 showed almost all increase due to i±2 interactions, while PN1 also showed notable increase at i±3, i±4, and trans contacts. Contacts at i±1 did not change substantially for either samples indicating that while the closely juxtaposed nucleosomes persist in the crosslinked material, no additional folding due to chromatin condensation involves interactions between the nearest neighbor nucleosomes.

Remarkably, the overall contact pattern in PN1 closely resembles that previously observed by EMANIC in interphase HeLa cells, while the one in PN56 is rather similar to reconstituted clone 601 chromatin showing strong traits of the 2-start zigzag (64). It thus appears that a major structural transition leading to the tight chromatin packing in mature retina originates from a folding according to the two-start zigzag despite the apparent absence of regular and helical 30 nm fibers.

### 7. Mesoscale chromatin modeling predicts that the short and irregular linker length distributions typical of PN1 compromise chromatin compaction

Mesoscale chromatin modeling at nucleosome resolution can assess multiple nucleosome chain configurations to detect histone tail-mediated interactions and other conformational details of internucleosomal interaction patterns and chromatin compaction (67,68). Here we developed two chromatin models for two selected PN1 and PN56 arrays with 23-nucleosomes (Fig. 7 A) to investigate the relationship between linker length distributions and chromatin condensation. For chromatin folding experiments, we modeled chromatin fibers with fully saturated linker histone (1 LH per nucleosome) and simulated chromatin folding at 5 mM, 50 mM or 150 mM NaCl, as well as in the presence or absence of 1 mM MgCl_2_.

**Figure 7.**
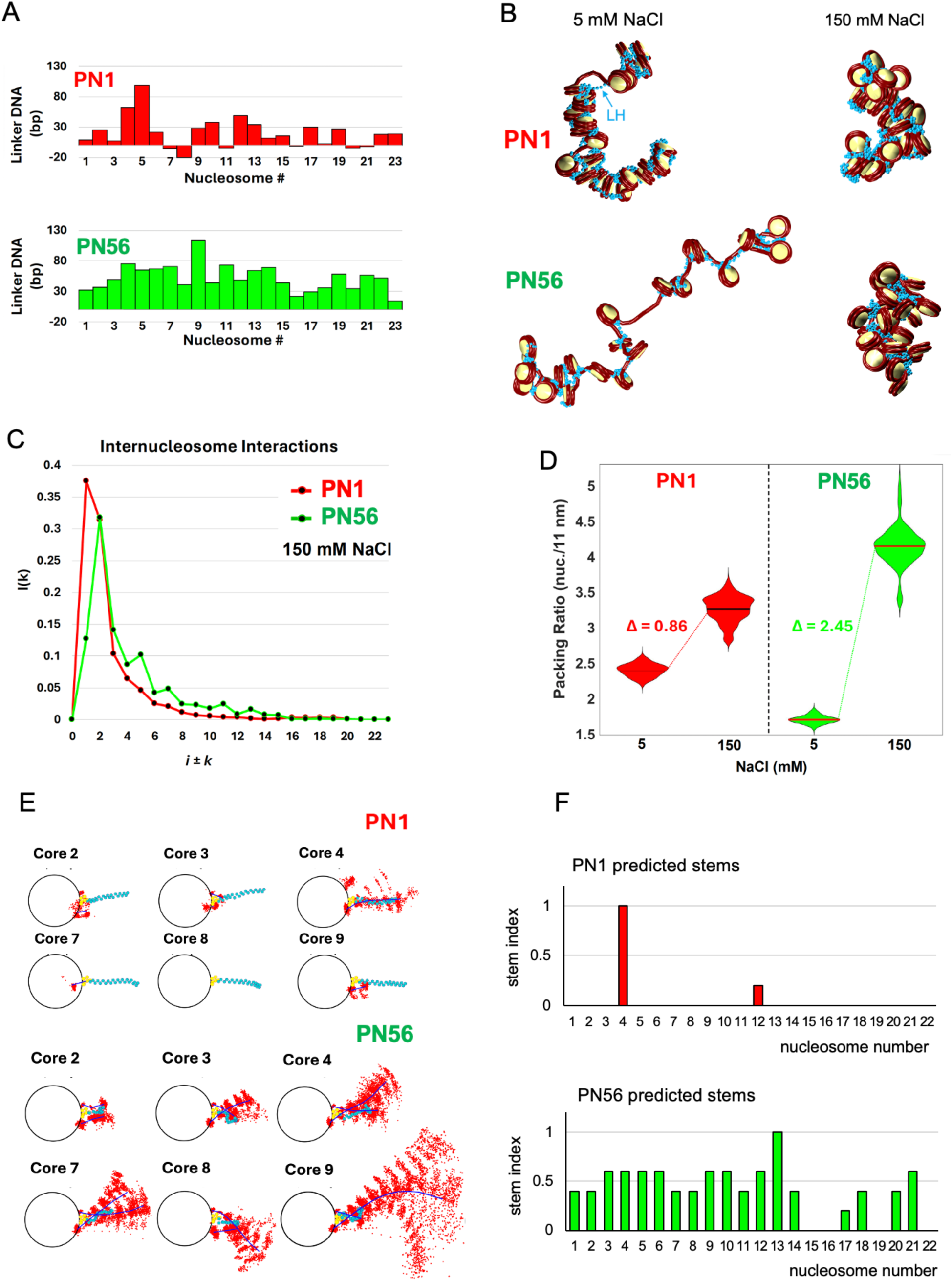
Mesoscale chromatin modeling and structural analysis of native chromatin from immature and mature retina. A: Bar graphs showing linker length L measured for a PN1 nucleosome array (red, CAP 1_4), and PN56 array (green, CAP 9_2). B: Representative conformations are shown for PN1 nucleosome arrays (top) and PN56 arrays (bottom) simulated at monovalent salt concentration of 5 and 150 mM NaCl as indicated. C: Nucleosome interaction patterns *i ± k* are shown for PN1 model (red) and PN56 model (green) with 1 linker histone per nucleosome at monovalent salt concentration of [NaCl] = 150mM. D: Chromatin packing ratios measured as the number of nucleosomes per 11 nm of the main fiber axis are shown for PN1 nucleosome arrays (red) and PN56 arrays (green) simulated at monovalent salt concentration of 5 and 150 mM NaCl as indicated. E: Fan plots representing, the linker DNA cumulative and average positional distribution across a single trajectory along each nucleosomal plane for selected PN1 and PN56 nucleosomes with shorter linkers (in PN1) and longer linkers (in PN56) modeled with 1 linker histone per nucleosome at monovalent salt concentration of [NaCl] = 150 mM. F: Bar graphs showing distributions of the predicted nucleosome stem formation index for the PN1 and PN56 nucleosome arrays modeled with 1 linker histone per nucleosome at monovalent salt concentration of [NaCl] = 150 mM.

As shown on Fig. 7 B-D, at physiological salt the two types of fibers predicted for PN1 and PN56 chromatin strikingly recapitulate the experiments: predominantly i±1 interactions in PN1 versus i±2 contacts in more compact PN56 (Fig. 7 B, C). Remarkably, at low salt, the PN56 array appears to be much less compact than PN1, mostly due to disordered long linkers. At physiological salt the long linkers fold compactly in the PN56 fiber, exhibiting a much stronger total amplitude for the salt-dependent chromatin packing measured as the number of nucleosomes per 11 nm unit length (Fig. 7 B, D). Our modeling is thus consistent with experimental observation of the much higher amplitudes of PN56 nuclear chromatin compaction (Fig. 1 C) and chromatin compaction by *in-situ* crosslinking (Fig. 6 F, H), and helps detect the underlying mechanism.

Namely, by plotting the configurations of the nucleosome linkers exiting and entering the nucleosomes with different linker length (Fig. 7 E, F), we see that the stem formation depends critically on linker DNA length. For nucleosomes with both linkers longer than 26 bp linker DNA stems form (narrow angle *α*) promoting the i±2 contacts between the neighboring nucleosomes. In contrast, shorter linkers cannot form stems (Fig. 7 F) and therefore reduce i±2 interactions and decrease overall fiber compaction for PN1 at physiological salt (Fig. 7 B). From tail interaction analysis of PN1 versus PN56 systems at two salt conditions, we also suggest that PN1 with short linkers has less tail-DNA interactions but more tail-tail and tail-non parental core interactions than the PN56 system of longer linkers. Thus, the high frequency of short linkers critically inhibits nucleosome chain folding and may explain the unfolded chromatin states observed in immature retina and possibly other proliferating cells. These features are in dramatic contrast with more ordered zigzag in mature chromatin.

## Discussion

One of the most intriguing natural biological phenomena associated with cell differentiation and tissue development involves accumulation of abundant facultative heterochromatin and its spatial segregation from euchromatin (83,84). Mammalian retina is a part of the central nervous system that has been studied extensively in relation to general epigenetic and chromatin-mediated mechanisms of neuronal differentiation (44–46) and specific mechanisms of retina photoreceptor reprogramming for treatment of blindness and improving vision (47–49).

Initially, based on the previous observation of 30-nm structural signal in mature retina and experiments with reconstituted chromatin with various NRLs (22,69,85), we expected that the increased NRL and levels of histone H1 in mature retina (45) would lead to formation of condensed 30-nm chromatin fibers while the shorter nucleosome repeat would promote an open nucleosome zigzag. However, our Cryo-ET analysis revealed remarkable structural heterogeneity incompatible with any helical structures in condensed chromatin of either immature or mature retina (Fig. 2). With open nucleosome arrays, the linker DNA length is strikingly irregular in the immature retina and deviates from any kind of regular structure. We propose that in the nucleus of immature retina cells, the open chromatin is maintained by the very frequent short linkers that effectively inhibit compact nucleosome folding (Fig. 8, left). Additionally, external nuclear structures such as ribonucleoprotein scaffolds (86) support the open chromatin state while the nuclear lamina (52) tethers heterochromatin at the nuclear periphery. During retina maturation, nucleosomes reorganize to acquire longer linker DNA lengths (Fig. 4 D) that promote formation of discontinuous 2-start zigzag structures and their accordion-like compaction (Fig. 8, right). At this stage, the attachments between heterochromatin and the nuclear lamina are disrupted allowing heterochromatin to congregate near the center of the nucleus (52). It thus appears that during retina cell maturation and chromatin inversion there is no abrupt transition from irregular chromatin to regular 30-nm fibers. Rather, the increasing linker DNA length and decreasing fraction of overlapping nucleosomes allow the chromatin to gradually increase the two-start zigzag features and fold more compactly, despite residual short linkers that interrupt the continuity of the chromatin fiber.

**Figure 8.**
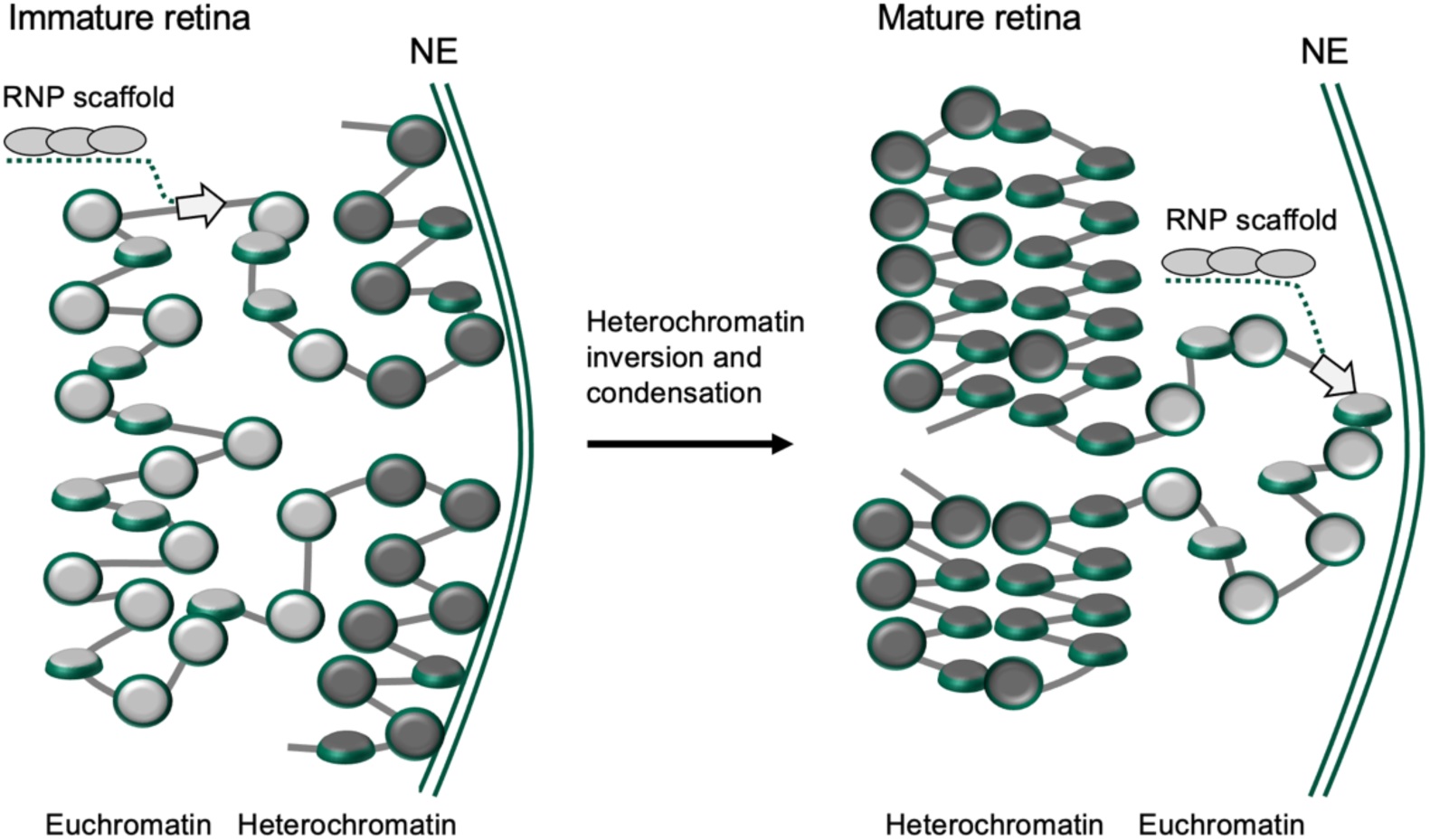
Schematic model of chromatin higher-order folding controlled by nucleosome spacing in naturing mouse retina. Schematic drawing of nucleosome chain folding in immature (left) and mature (right) mouse retina cells chromatin implies that that the closely spaced di- and tri-nucleosomes render nucleosome arrays more open in immature rod photoreceptors. During retina maturation, heterochromatin fraction increases, number of closely-spaced nucleosomes decreases, and the chromatin fibers become globally condensed via tight accordion-like zigzag folding. This mechanism may be a general way of maintaining a more open chromatin to ensure high epigenetic plasticity in immature and undifferentiated cells and specialized gene expression programs in differentiated and mature cells. NE: nuclear envelope.

Remarkably, a significant fraction of nucleosomes in immature retina chromatin has negative linker length values (e.g., L7, L8, and L11 on Fig. 4 B) and thus contains partially overlapping two or three nucleosome cores. Such overlapping nucleosomes were initially observed as a result of chromatin remodeling by SWI/SNF *in vitro* (87). With native chromatin, overlapping nucleosomes were observed by Cryo-ET in human HeLa chromosomes (60) and Drosophila embryos (88). By genome-wide mapping overlapping nucleosomes were found to be enriched at transcriptional start sites in human HeLa cells (89,90) and actively transcribed genes in murine embryonic stem cells (91). Structures of overlapping di- and tri-nucleosomes reconstituted from nucleosome positioning DNA and mixture of histone octamers and hexamers lacking one histone H2A/H2B dimer have been solved by X-ray crystallography (89) and Cryo-EM (90). Another interesting example of partially overlapping nucleosomes has been reconstituted *in vitro* from telomeric DNA repeats and analyzed by cryo-EM (40).

Cryo-ET analysis of mouse retina chromatin reveals several new features of the partially overlapping nucleosomes. First – they are abundant in immature retina and their frequency (10.7 % of total linkers in PN1) appears to be much higher than that expected from transcription start sites. Second, the extent of the nucleosome overlap is variable (Fig. 4 E) and the strictly overlapping nucleosomes (with negative-length linkers) do not make a structurally homogeneous unit but rather include a group of closely juxtaposed nucleosomes with very short linkers. Third, the overlapping nucleosomes can be also found in the mature retina where they are less abundant and often interspersed, within the same array, with the longer-linker nucleosomes containing stems.

The unprecedented heterogeneity of linker DNA length makes the native chromatin folding incompatible with previous nucleosome models based on regular helical nucleosome arrangements and average NRL (43). Here, to account for the native nucleosome positional heterogeneity, we generated for the first time nucleosome-resolution chromatin models with individual nucleosome linker length variations derived from native mouse retina chromatin. Our simulated chromatin fiber systems of PN1 and PN56, remarkably consistent with experiments, suggest a new mechanism: partially overlapped nucleosomes are sufficient to perturb chromatin higher-order folding in immature mouse retina. Interestingly, not just the overlapped nucleosomes, but all nucleosomes with rather short linkers, below 26 bp, limit formation of the nucleosome stem and thus impede histone H1-induced chromatin folding (**Fig. 7**). That the length of the linker DNA perturbing stem formation (< 26 bp) is very close to those constrained by histone H1 positioned near the dyad axis of a chromatosome (42,92) and directly contacting histone H1 C-tail domain (93) is consistent with perturbance of stem formation being the main mechanism by which the short linkers inhibit chromatin folding.

What are the factors that regulate nucleosome spacing and cause partial nucleosome overlapping in immature retina? And what leads to the increased nucleosome spacing upon retina cell maturation? In living cells, nucleosome spacing is dynamically regulated by ATP-dependent chromatin remodelers. (94). As we discussed above, the overlapped nucleosomes result from the activity of nucleosome remodelers SWI/SNF *in vitro* and are attenuated by murine SWI/SNF homolog BRG1 (Smarca4) and ISWI homolog SNF2H (Smarca5) in mouse embryonic stem cells (91). Remarkably, Smarca4 is important for cell integrity in early retina progenitors (95) and SmarcA5 is essential for retinal cell proliferation and photoreceptor maintenance (96). Another factor essential for maintenance of the NRL is histone H1 (97) whose expression is increased in mature mouse retina and its partial knockout promotes antigenic exposure but not unfolding of the facultative heterochromatin (45). Whether the chromatin remodelers and/or histone H1 levels control the number of overlapped nucleosomes in developing retina remains to be determined.

Together, the observation of abundant overlapping nucleosomes due to short and irregular nucleosome linkers in immature retina, with development-mimicking chromatin models predicting compromised chromatin folding, define a new mechanism for developmentally-regulated control of chromatin structure in mouse retina. The precise nature of nucleosome positioning (or position-randomizing) factors and how such mechanisms affect chromatin compaction in other cells and tissues are exiting areas for future research. Indeed, for mouse ES cells, HeLa, and Drosophila embryos where the partially overlapped nucleosomes have been associated with open chromatin, new mechanistic questions on the role of nucleosome distribution in the chromatin folding plasticity and epigenetic regulation naturally emerge.

## Acknowledgements

We are thankful to J. Sloppy for technical assistance with electron microscopy at the Penn State Hershey Cryo-EM Facility (Facility SCR_021178) and SURIP program undergraduate students C. Rohrer, K. Chowdhurry, and O. Plotts for research assistance.

